# Interoperability of RTN1A in dendrite dynamics and immune functions in human Langerhans cells

**DOI:** 10.1101/2022.04.08.487626

**Authors:** Małgorzata Anna Cichoń, Karin Pfisterer, Judith Leitner, Lena Wagner, Clement Staud, Peter Steinberger, Adelheid Elbe-Bürger

**Affiliations:** Department of Dermatology, Medical University of Vienna, Austria; Center for Pathophysiology, Infectiology and Immunology, Medical University of Vienna, Austria; Department of Plastic and Reconstructive Surgery, Medical University of Vienna, Austria

## Abstract

Skin is an active immune organ where professional antigen-presenting cells such as epidermal Langerhans cells (LCs) link innate and adaptive immune responses. While Reticulon 1A (RTN1A) was recently identified in LCs and dendritic cells in cutaneous and lymphoid tissues of humans and mice, its function is still unclear. Here, we studied the involvement of this protein in cytoskeletal remodeling and immune responses towards pathogens by stimulation of Toll-like receptors (TLRs) in resident LCs (rLCs) and emigrated LCs (eLCs) in human epidermis ex vivo and in a transgenic THP-1 RTN1A^+^ cell line. Hampering RTN1A functionality through an inhibitory antibody induced significant dendrite retraction of rLCs and inhibited their emigration. Similarly, expression of RTN1A in THP-1 cells significantly altered their morphology, enhanced aggregation potential and inhibited the Ca^2+^ flux. Differentiated THP-1 RTN1A^+^ macrophages exhibited long cell protrusions and a larger cell body size in comparison to wild type cells. Further, stimulation of epidermal sheets with bacterial lipoproteins (TLR1/2 and TLR2) and single-stranded RNA (TLR7) resulted in the formation of substantial clusters of rLCs and a significant decrease of RTN1A expression in eLCs. Together, our data indicate involvement of RTN1A in dendrite dynamics and structural plasticity of primary LCs. Moreover, we discovered a relation between activation of TLRs, clustering of LCs and downregulation of RTN1A within the epidermis, thus indicating an important role of RTN1A in LC residency and maintaining tissue homeostasis.

**Graphical abstract:** 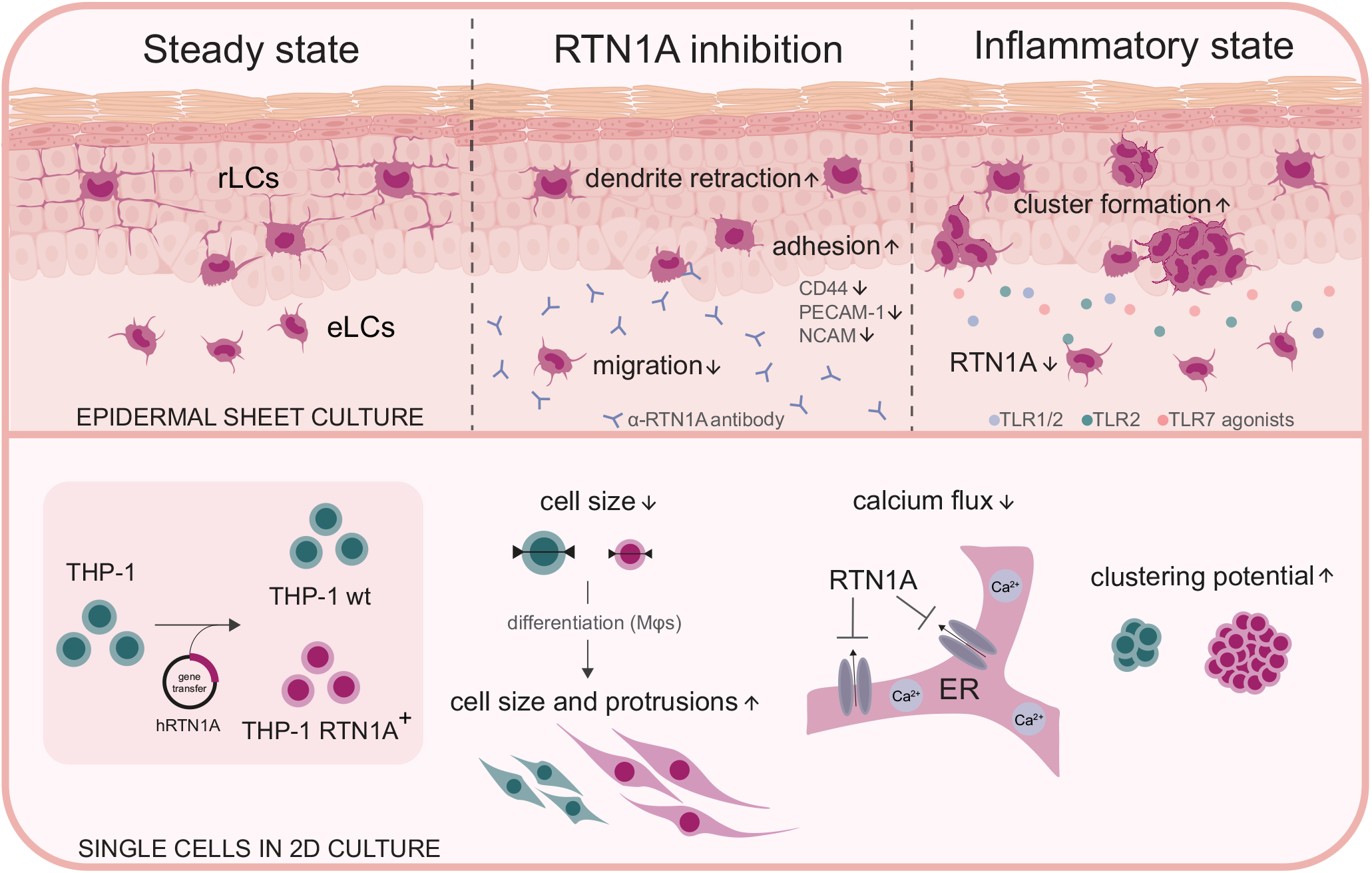

**Highlights:**

- Blocking of RTN1A induces dendrite retraction of resident LCs (rLCs) in epidermal explants.
- Despite a roundish morphology rLCs exhibit reduced migration capacity.
- RTN1A has an inhibitory effect on the calcium flux.
- Toll-like receptor-activated rLCs form vast clusters and significantly diminish RTN1A expression after emigration.
- RTN1A plays a central role in LC residency and maintaining tissue homeostasis.

## Introduction

A well-functioning network of dendritic cells (DCs) and macrophages (Mφs) in the upper layer of human skin is of great importance to defeat invading pathogens and to sustain the intact skin barrier. Langerhans cells (LCs) reside in the epidermis in an immature state, have features of both DCs and Mφs and can function in maintaining tolerance but also priming an immune response^1–3^. Recently, more light was shed on LC diversity and how the phenotype of LC subtypes can vary depending on the function they are dedicated to^4–6^. Moreover, in healthy skin the LC fate can be shaped by the environment^7–9^. Responsiveness to external intruders results in activation and maturation of certain LC subsets sensing antigens via activation of toll-like receptors (TLRs), a category of pattern recognition receptors that initiate the innate immune response. They not only recognize specific microbial particles termed pathogen-associated molecular patterns including lipopolysaccharides of gram-negative and lipoteichoic acid of gram-positive bacteria, and nucleic acids of viruses, but also endogenous damage-associated molecular pattern molecules. TLRs which recognize nucleic acids, reside in intracellular compartments to decrease the risk to encounter “self” nucleic acids, whereas cell surface TLRs largely recognize microbial membrane compartments and therefore do not require this protective strategy^10–16^. Activated LCs downregulate molecules necessary for tissue exit such as E-cadherin^17,18^, upregulate costimulatory molecules, secrete cytokines and undergo a complex cytoskeleton remodeling that enables LCs to acquire a roundish shape with short dendrites to efficiently relocate within and through the epidermis towards dermal lymphatic vessels and lastly lymph nodes^19^. The crosstalk between LC activation and morphological changes is not fully understood. LCs can be distinguished by their morphological plasticity, namely their readily dendrite extension during their residence and replenishment in the epidermis^20,21^ and changes in dendrite patterning resulting in dendrite retraction due to their intra-tissue migration capacity^22^. LC dendrites are membranous extensions, which in healthy skin exhibit orderly dendrite distribution with small distance to other cells^23^. Notably, dendrites can display similar structures, branching and intersection frequency as neurons^24^.

The endoplasmic reticulum (ER)-associated protein Reticulon 1A (RTN1A) was recently identified in DCs of cutaneous and lymphoid tissues in humans and mice^25,26^. Previously an interesting observation was made on the distribution of the RTN1A protein within LC dendrites^25^. Indeed, some ER-associated proteins (e.g. Atlastin-1), have been shown to be involved in the distribution and shaping ER morphology in neural dendrites^27^. Certain RTN-family members have been associated with ER morphogenesis^28,29^. In neuroanatomy research, the involvement of RTNs in neurite expansion and regeneration was well investigated. For example, RTN4 has been shown to be involved in inhibition of the neurite outgrowth in human cell lines^30^ and inhibition of RTN4A via an antibody (ab) can improve visual recovery after retinal injury in mice^31^. RTN1, in contrast to RTN4, did not inhibit axonal regeneration^32^, which correlates with our previous observations that RTN1A might be involved in axonal elongation of cutaneous nerves in prenatal mouse skin^25^. In humans, a possible interaction between the cytoskeleton and RTN4 in monocyte-derived Mφs for instance has been pointed out^33^. Based on previous findings, we aimed to add understanding on the function of RTN1A in the ER of LCs and study its involvement in structural remodeling. To identify the function and potential involvement of RTN1A-induced morphological changes we studied its behavior in immune responses to specific TLR agonists on/in resident LCs (rLCs) and emigrated LCs (eLCs) in human epidermal explants.

## Results

### RTN1A is involved in the dynamic of dendrite retraction in LCs

In the first set of experiments we studied morphological changes in rLCs upon hampering RTN1A functionality with an α-RTN1A ab in human epidermal sheets *ex vivo* (Fig 1A). Incubation of dermatomed skin with the enzyme dispase II dissociated the basement membrane, thus enabling separation of the epidermis from the dermis and consequently exposure of epidermal cells to abs. The untagged α-RTN1A ab was detected with a fluorescently-labelled secondary ab in the cell body and in the dendrites of rLCs already after only 3 h, more pronounced at 6 h and most prominent in the majority of rLCs at 24 h of cultivation, while the isotype signal was substantially weaker (6 h) or undetectable (3 and 24 h) (Fig 1B,C). A prominent roundish morphology of rLCs was observed after 24 h of incubation with the α-RTN1A abs in comparison to isotype-treated epidermis. Notably, only RTN1A^+^ rLCs captured the RTN1A ab (Fig 1C, yellow arrows). Indeed, not all rLCs and eLCs express RTN1A. We analyzed the frequency of RTN1A expression in human primary LCs and found that around 80% of rLCs and eLCs express this protein (S. Fig 1A), from which ∼70% of all rLCs and eLCs co-express CD207 and CD1a, respectively (S. Fig 1B,C). For further analysis of rLCs in this experimental set up we employed markers such as the C-type lectin receptor CD207 (can be localized in the plasma membrane and intracellularly)^22^, RTN1A (distributed in the ER) and vimentin to label intermediate filaments (major component of the cytoskeleton)^34^. Images of untreated (UT) epidermal sheets cultured for 24 h showed co-localization of RTN1A with vimentin in the cell body and in the most distant tips of dendrites with dot-like RTN1A accumulations (Fig 1D, pinkish arrows), whereas CD207 expression was detected at lower intensity and mainly the cell body (Fig 1D). We also captured cells with halfway-retracted dendrites most likely representing migratory LCs (Fig 1D, white arrows). Upon 24 h of incubation with α-RTN1A abs, rLCs revealed a consistent and recurrent reduction of dendricity in comparison to isotype-treated and UT epidermal sheets (Fig 1D). Enumeration of dendritic and roundish rLCs within epidermal biopsy punches after 24 h of incubation with or without abs showed a significant decrease in the number (Fig 1E) and percentage (UT: 71.47; isotype: 63.52; α-RTN1A ab: 39.74; Fig 1F) of dendritic rLCs after blocking of RTN1A in comparison to controls. This observation correlates with a significant increase of roundish rLCs (UT: 28.53; isotype: 36.48; α-RTN1A ab: 60.26; Fig 1F).

**Fig 1.**
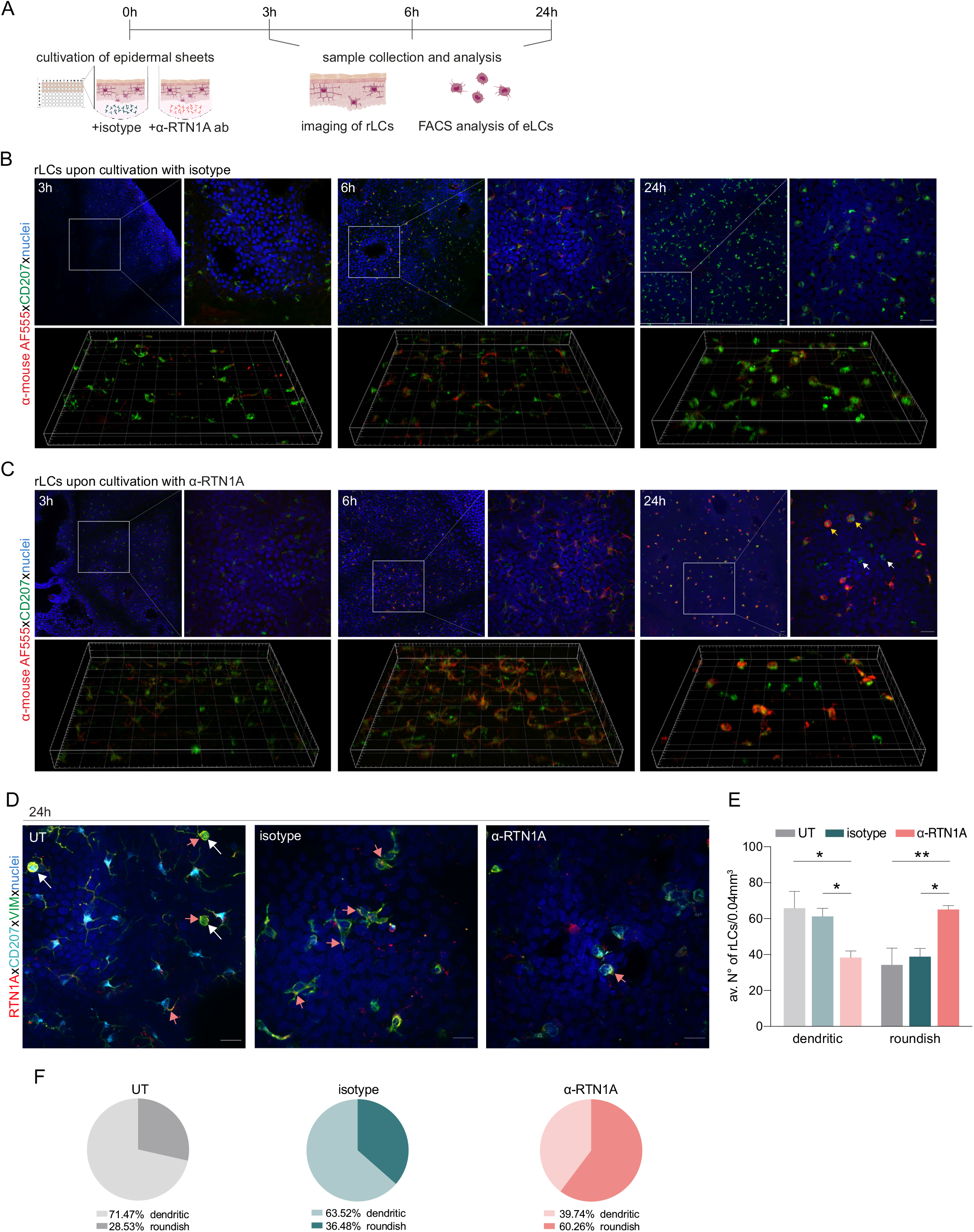
Impairment of RTN1A functionality instigates a roundish morphology in resident Langerhans cells (rLCs). A. Experimental workflow demonstrating the cultivation of human epidermal sheets with an α-RTN1A ab and the isotype for indicated time points and subsequent analysis strategies. B-C. Representative immunofluorescence (IF) images of isotype- and α-RTN1A-treated epidermal sheets at indicated time points stained for CD207 (green), a secondary ab (red) to visualize the uptake of the isotype and α-RTN1A ab by rLCs and nuclear counterstaining with DAPI (blue). Zoom-ins of the boxed areas are also shown as 3D projections underneath. RTN1A^+^ rLCs: yellow arrows, RTN1A^-^ rLCs: white arrows. n=3, scale bar: 20 μm. D. Representative IF images showing RTN1A, CD207, vimentin and DAPI staining in untreated (UT), isotype- and α-RTN1A-treated human epidermal sheets after 24 h of cultivation. Colocalization of RTN1A with vimentin: pinkish arrows, rLCs with partially retracted dendrites: white arrows. n=4, scale bar: 20 μm. E, F. Enumeration, percentage and distribution of dendritic and roundish rLCs in epidermal sheets upon 24 h of culture and indicated treatment. E, Data are shown as standard error of the mean (SEM) from 4 fields of view (FOV) of 4 donors and were analyzed using two-way ANOVA with Tukey’s multiple-comparison test. F, Data represent mean of 4 donors. \**p* ≤ 0.05, \*\**p* ≤ 0.01

Next, we compared morphological changes in rLCs in UT, isotype and α-RTN1A ab-treated epidermal sheets. Using 3D trajectories of intermediate filaments (vimentin) in rLCs (Fig 2A), we analyzed cell dendricity and the frequency of dendrite distribution in rLCs (Fig 2B,C). Quantification of the dendrite lengths revealed that inhibition of RTN1A caused significant dendrite retraction in rLCs compared with those in isotype and UT epidermal sheets (Fig 2B). Sholl analysis addressing frequency and complexity of dendrite brunches showed significantly decreased rLC dendrite distribution in α-RTN1A ab-treated epidermal sheets in comparison to controls and most pronounced to UT epidermal sheets (Fig 2C).

**Fig 2.**
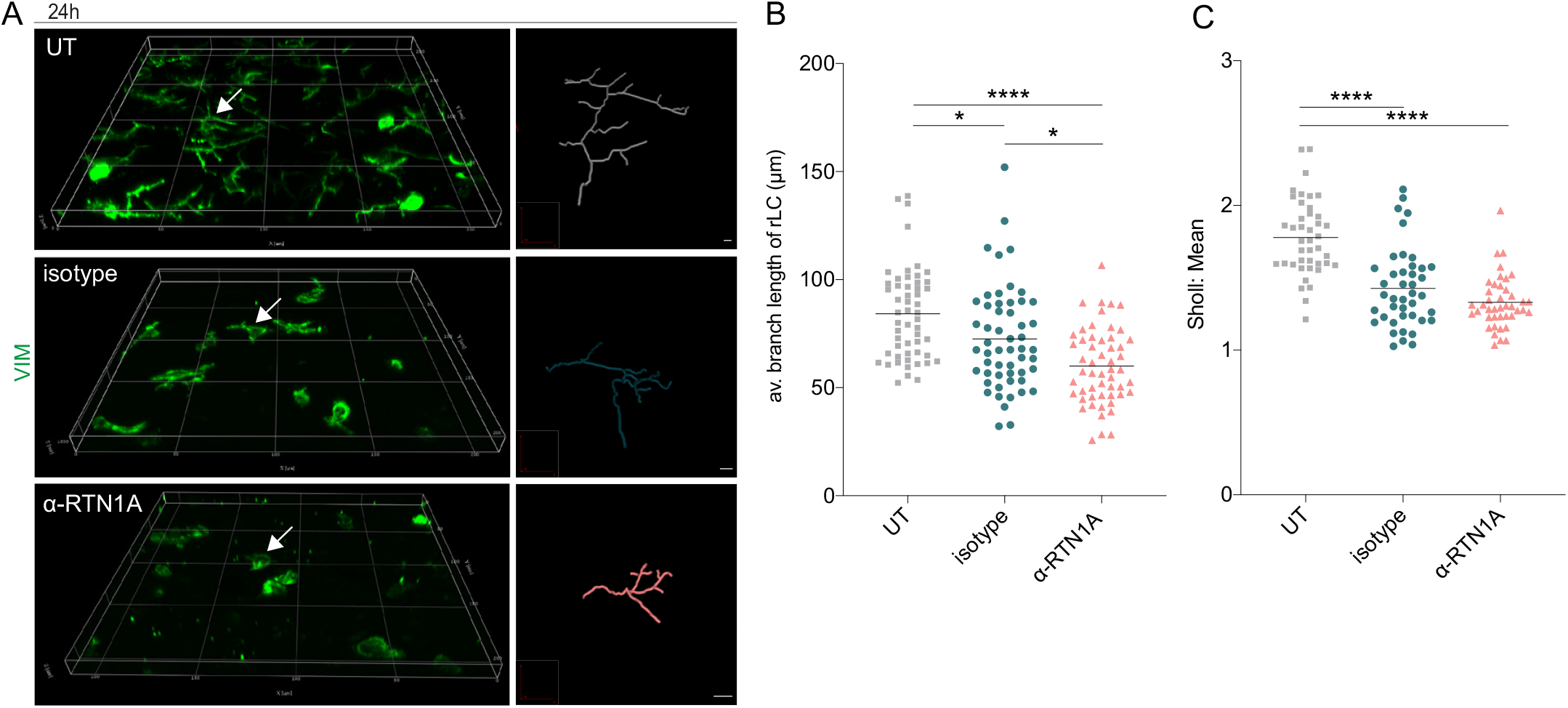
Inhibition of RTN1A in resident Langerhans cells (rLCs) significantly alters dendrite length and distribution. A. 3D projections of epidermal sheets upon 24 h of culture, indicated treatment and vimentin (VIM) staining (left panel). Shown are single rLC 3D trajectories based on intermediate filament expression (right panel). Scale bar: 10 μm. B, C. Evaluation of the average rLC branch lengths and Sholl analysis of rLC dendricity upon 24 h of culture and indicated treatments. 4 FOV were evaluated of 4 donors and data analyzed using two-way ANOVA with Tukey’s multiple-comparison test, \**p* ≤ 0.05, \*\*\*\**p* ≤ 0.0001.

### Despite dendrite retraction, rLCs with blocked RTN1A function remain in the epidermis

To evaluate whether blocking of RTN1A with the inhibitory ab affects LC emigration from epidermal sheets, we analyzed the number of eLCsat selected time-points. We found a significant decrease in the migration potential of LCs in culture wells containing α-RTN1A ab-treated compared to isotype-treated epidermal sheets after 24 h of cultivation (Fig 3A). Similar to rLCs (Fig 1B,C), ab uptake was also detected in eLCs at all investigated time points, with a tendency of a more pronounced though not significantly higher signal of RTN1A ab compared to the isotype (Fig 3B). We assume that LCs carried the α-RTN1A/isotype abs after detachment from the epidermis. However, it is also conceivable that some eLCs have taken up abs in the medium after the emigration (Fig 3B). Next, we analyzed typical LC markers in eLCs (Fig 3C) such as CD1a, a microbial lipid-presenting molecule^35,36,22^, as well as CD207 We found that the CD1a expression level (mean fluorescence intensity; MFI), but not the percentage of CD1a^+^ eLCs was significantly reduced after α-RTN1A ab treatment, whereas CD207 expression was unchanged. The migratory and mature phenotype of LCs is distinguishable by increased expression levels of CCR7 and co-stimulatory molecules such as CD86^37^. Here, the inhibition of RTN1A caused a significant increase in the percentage and expression intensity of CCR7, and a decrease of CD86 in comparison to controls. These results suggest that inhibition of RTN1A endorse acquisition of the migratory phenotype by LCs, along with preventing the maturation of some LCs. To better understand the mechanism involved in rLC morphology changes after blocking RTN1A functionality and LC migration, we comparatively measured the presence of key adhesion molecules in culture supernatants after 24 h. The adhesion molecule CD44 can stimulate intracellular calcium mobilization, and actin-^38^ and vimentin-mediated cytoskeleton remodeling^39^. PECAM-1^40^ plays a role in endothelial cell-cell adhesion^41^ and NCAM promotes neuron-neuron adhesion and neurite outgrowth^42^. ALCAM facilitates attachment of DCs to endothelial cells^43^ and migration of other endothelial cells^44^, EpCAM regulates LC adhesion and foster their migration^45,46^. L-selectin is a calcium-dependent lectin expressed by leukocytes and mediates cell adhesion by binding to neighboring cells^47,48^. Treatment of epidermal sheets with the α-RTN1A ab for 24 h, significantly decreased the concentrations of CD44, PECAM-1, NCAM, ALCAM and EpCAM but not L-selectin in culture supernatants in comparison with controls (Fig 3D). These data suggest that the detachment of rLCs from the tissue was hampered.

**Fig 3.**
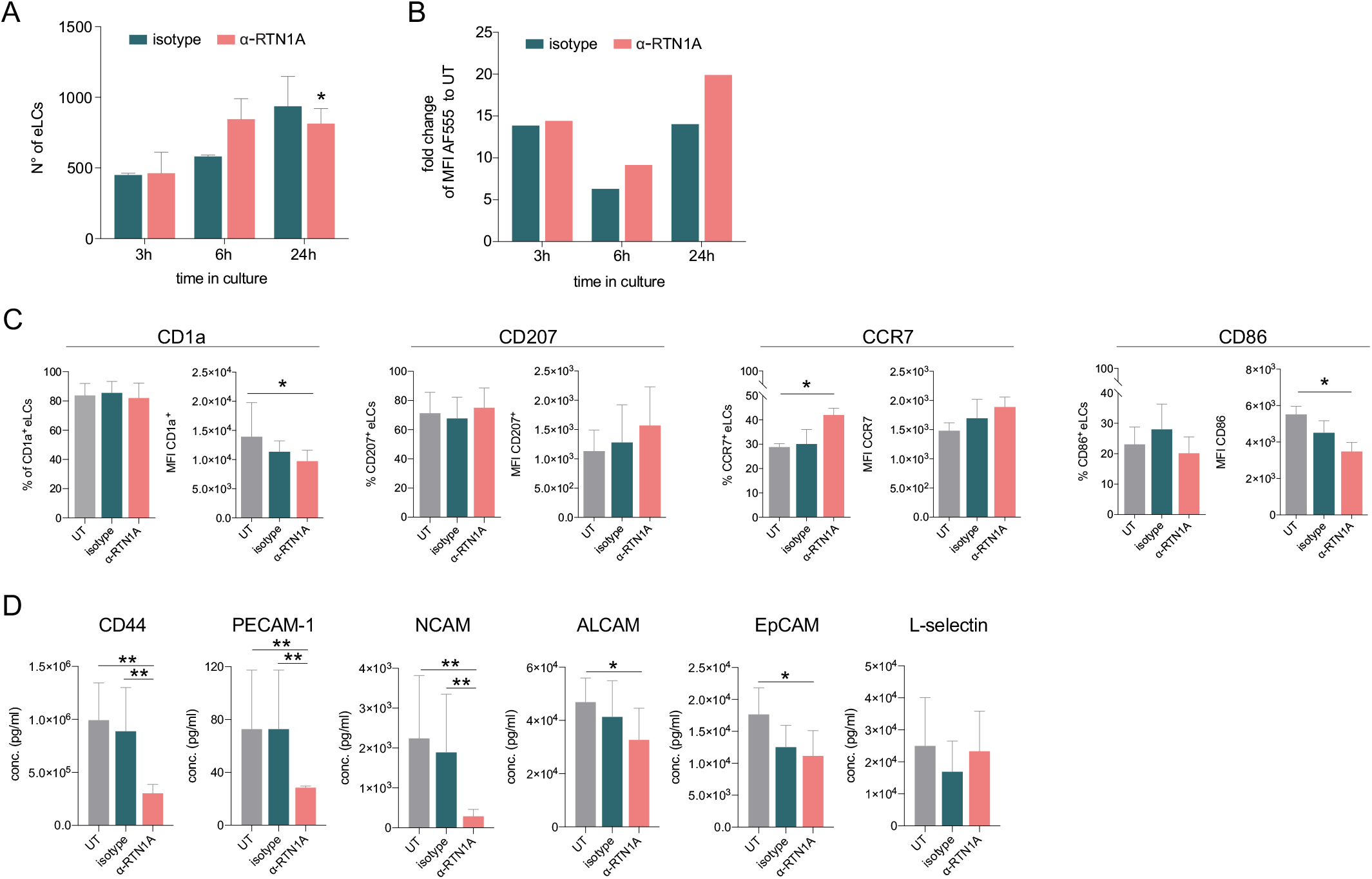
Hampering RTN1A function decreases the Langerhans cell (LC) migration potential and alters the marker expression in emigrated LCs (eLCs). A, B. Enumeration and ab uptake of emigrated LCs (eLCs), collected from culture wells with epidermal sheets at indicated time points and analyzed via flow cytometry. A, Data are presented as standard error of the mean (SEM) from triplicates of 3 donors. Ordinary one-way ANOVA with Tukey’s multi-comparison test was used. \**p* ≤ 0.05. B, Data from triplicates, including 3 donors are shown as mean fold change to untreated (UT) eLCs. C. Marker expression profile of eLCs upon 24 h of culture, treatment, and flow cytometry analysis. Data shown represent mean ± SEM of 3 donors and were analyzed using two-way ANOVA Tukey’s multi-comparison test. \**p* ≤ 0.05 D. Adhesion molecule concentrations in supernatants of cultivated epidermal sheets with indicated 24 h treatments and subsequent LEGENDplex bead array measurement. Data are shown as SEM from duplicates of 4 donors and were analyzed using two-way ANOVA with Tukey’s multiple-comparison test. \**p* ≤ 0.05, \*\**p* ≤ 0.01

### Expression of RTN1A substantially changes the cell size of myeloid cells

Next, we assessed the involvement of RTN1A in cytoskeletal remodeling by determining shape and morphology changes upon expression of human RTN1A in the RTN1A^-^ monocyte-like cell line THP-1 (Fig 4A). After confirming protein expression by flow cytometry, THP-1 RTN1A^+^ cells and THP-1 wild type (wt) cells were used for further experiments (Fig 4A). THP-1 RTN1A^+^ cell morphology was considerably altered compared to THP-1 wt cells as they were significantly smaller in size and had markedly more condensed intermediate filament structures as visualized by vimentin staining (Fig 4B,C and S. Fig 2A). To assess a potential cross-talk between RTN1A and the cytoskeleton, THP-1 RTN1A^+^ cells were cultivated on fibronectin and imaged to analyze co-localization of RTN1A with vimentin and F-actin in three cell compartments (bottom, middle and top) (Fig 4D, S. Fig 2B). We found a significant overlap between RTN1A and vimentin at the bottom and top of the cell (Fig 4D,E), whereas there was less colocalization with F-actin in the same cell compartments (Fig 4D,E). In the middle part of the cell, RTN1A showed similar colocalization levels with F-actin and vimentin (Fig 4D,E). As during cultivation THP-1 wt and THP-1 RTN1A^+^ cells showed different growth dynamics, we subsequently evaluated their proliferation rate and cell growth.

**Fig 4.**
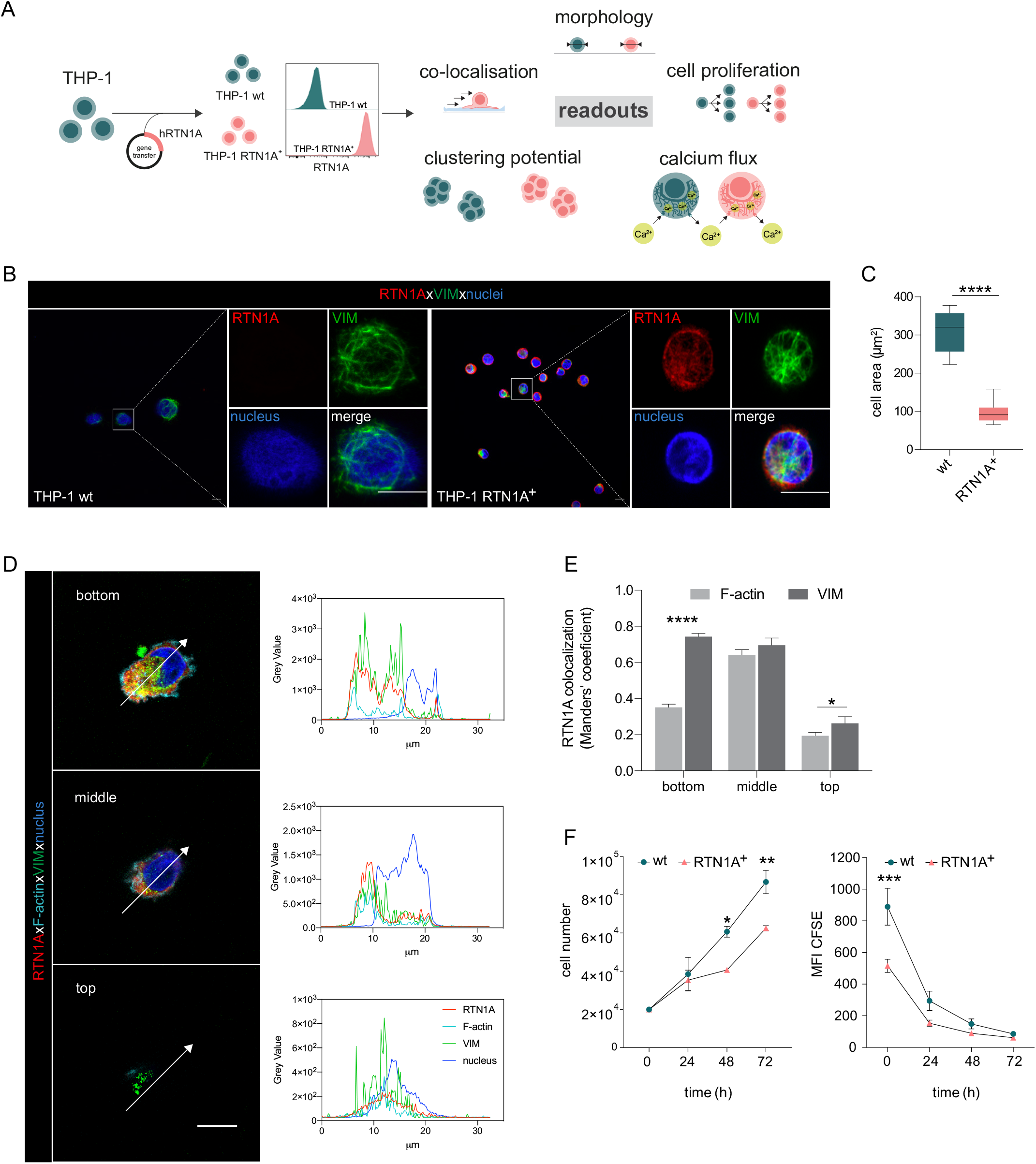
Expression of RTN1A in the myeloid THP-1 cell line affects the cell size. A. Workflow for the generation of THP-1 RTN1A^+^ cells and their comparative analysis with THP-1 wild type (wt) cells. B. Representative immunofluorescence (IF) images of THP-1 wt and THP-1 RTN1A^+^ cells on adhesion slides stained for RTN1A, vimentin (VIM) and nuclei (DAPI). n=4; scale bar: 10 μm. C. Comparative evaluation of the cell area revealed substantial divergences between THP-1 wt and THP-1 RTN1A^+^ cells. Data are shown as standard error of the mean (SEM) from 4 fields of view (FOV; n=4) and were analyzed using unpaired, two-tailed student t test. \*\*\*\**p* ≤ 0.0001 D, E. Representative IF images and quantification using Manders’ coefficient of RTN1A colocalization with filamentous proteins in a THP-1 RTN1A^+^ cell within 3 cell compartments: bottom, middle and top of the cell (right panel). Scale bar: 10 μm. Data are shown as SEM (10 cells/4 FOV; n=2), and analyzed using two-way ANOVA with Tukey’s multiple-comparison test. \**p* ≤ 0.05, \*\*\*\**p* ≤ 0.0001 F. Evaluation of the cell number and proliferation rate of THP-1 wt and THP-1 RTN1A^+^ cells within the time period indicated. Data presented as SEM were analyzed with two-way ANOVA, Sidak’s multiple comparison test (n=3). \**p* < 0.05, \*\**p* ≤ 0.01, \*\*\**p* ≤ 0.001

Indeed, THP-1 RTN1A^+^ cells displayed a significantly lower proliferation rate, CFSE dye dilution and cell number compared with THP-1 wt control (Fig 4F). The involvement of RTN1A in determination of morphological features such as cell size and dendricity was further studied in differentiated adherent THP-1 RTN1A^-^ wt Mφs and THP-1 RTN1A^+^ Mφs (Fig 5A,B). Expression of RTN1A in THP-1 Mφs resulted in altered morphology with significantly larger cell bodies and substantially longer cell protrusions when compared with THP-1 wt Mφs (Fig 5B-D). Alike in undifferentiated THP-1 cells (Fig 4D,E), RTN1A significantly co-localized with vimentin and to a lesser extent with F-actin (Fig 5E),

**Fig 5.**
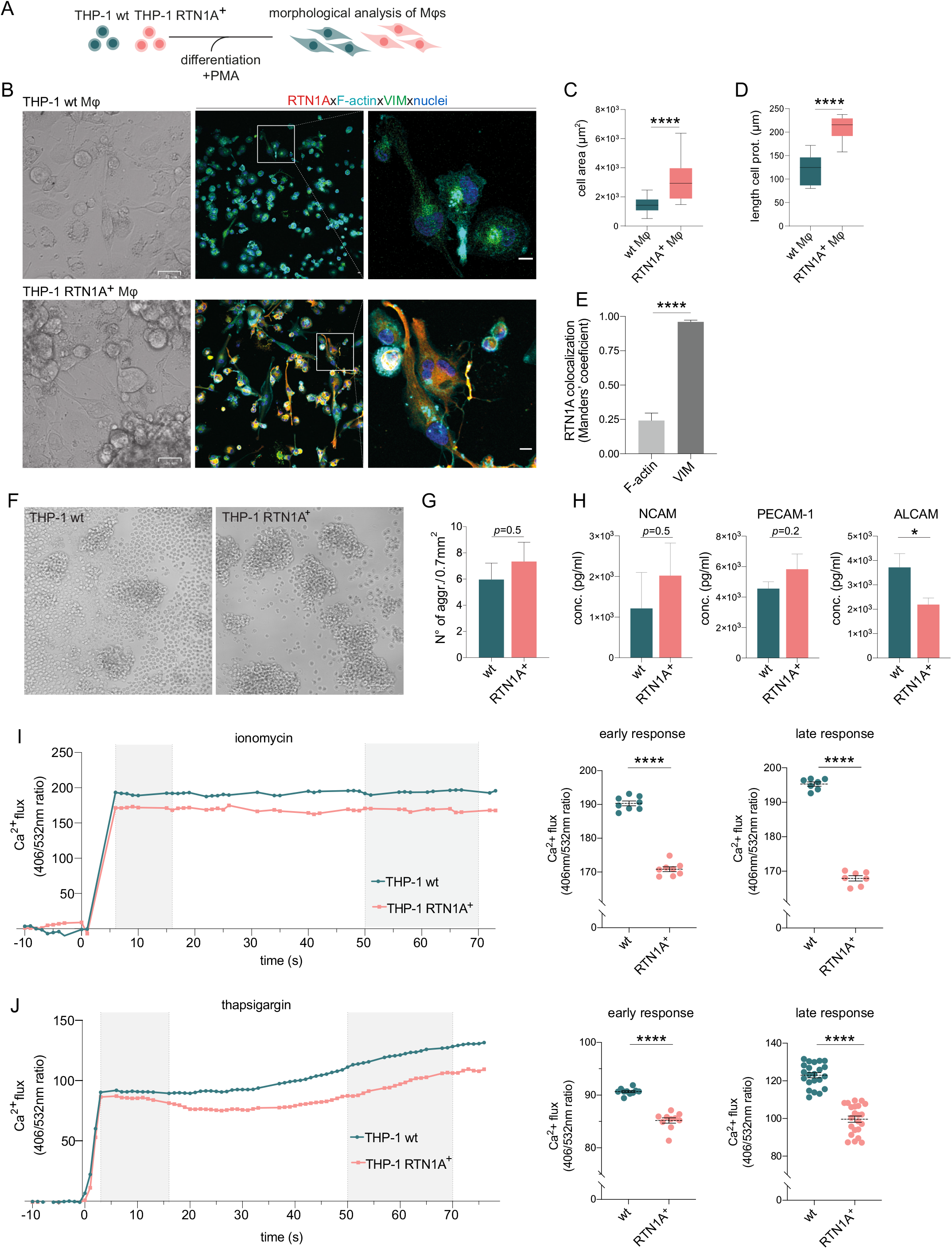
RTN1A considerably alters the morphology of THP-1 RTN1A^+^ Mφs as well as enhances aggregate formation and impairs Ca^2+^ flux in THP-1 RTN1A^+^ cells. A. Workflow for the differentiation and comparative analysis of THP-1 RTN1A^+^ Mφs and THP-1 wt Mφs. B. Representative bright field (BF) and immunofluorescence images (IF; co-staining with RTN1A, F-actin, vimentin (VIM), and DAPI) of THP-1 wt Mφs and THP-1 RTN1A^+^ Mφs. Scale bar: BF: 31 and IF: 10 μm, n=4. C, D. Comparative analysis of cell body size and average length of cell protrusions of differentiated THP-1 wt Mφs and THP-1 RTN1A^+^ Mφs. Data were analyzed using unpaired, two-tailed student t test, n=4. \*\*\*\**p* ≤ 0.0001 E. Comparative colocalization between RTN1A, F-actin and VIM using Manders’ coefficient. Data shown are standard error of the mean (SEM), n=4. \*\*\*\**p* ≤ 0.0001 F. Representative BF image of THP-1 wt and THP-1 RTN1A^+^ cells forming aggregates during culture. Scale bar: 100 µm. G. Enumeration of THP-1 wt and THP-1 RTN1A^+^ cell aggregates. Data are demonstrated as SEM from 4 fields of view per passage and analyzed using unpaired, two-tailed student t test, n=6. H. Adhesion molecule concentrations in supernatants after 48 h of THP-1 wt and THP-1 RTN1A^+^ cell cultivation, measured in duplicates with LEGENDplex bead array. Data are shown as SEM from 3 different passages and were analyzed using unpaired, two-tailed student t test. \**p* ≤ 0.05 I, J. Ratiomeric analysis of early and late phase calcium flux in THP-1 wt and THP-1 RTN1A^+^ cells using Fura-3 calcium indicator, ionomycin (n=5) and thapsigargin (n=4). Unpaired two-tailed t test was used for the response to both ionomycin and thapsigargin. \*\*\*\**p* ≤ 0.0001

### RTN1A inhibits calcium flux and regulates cell adhesion in myeloid cells

We discovered that THP-1 RTN1A^+^ cells display an increased capacity in cellular aggregate formation compared with THP-1 wt cells (Fig 5F,G and S. Fig 2C). In line with this observation, we analysed cell culture supernatants for the presence of adhesion molecules. We found increased NCAM and PECAM-1 and significantly decreased ALCAM concentrations in supernatants of THP-1 RTN1A^+^ in comparison to THP-1 wt cells (Fig 5H). The ER is an important calcium ion store and proper calcium levels are crucial for maintaining balanced cell functions^49,4^. Calcium ions (Ca^2+^) are a versatile second messenger involved in signal transduction and controlling activity of adhesion molecules, such as CD44, L-selectin^50,51^. To analyse whether an altered calcium homeostasis in THP-1 RTN1A^+^ cells promotes the enhanced aggregation capacity, we comparatively monitored Ca^2+^ flux in THP-1 wt and THP-1 RTN1A^+^ cells using ratiomeric calcium flux measurement^52^ with the Ca^2+^ indicator Fura-3 red. Calcium ionophores such as ionomycin are binding calcium ions^53^, which induces opening calcium stores and reaugmenting of [Ca^+^]i^54^. Both, the early and late response to ionomycin in THP-1 RTN1A^+^ cells was significantly lower in comparison to the calcium flux in THP-1 wt cells (Fig. 5I). Next, we tested whether RTN1A plays a role in differently induced Ca^2+^ mobilization from the ER and therefore applied thapsigargin^53^. This compound inhibits sarco/endoplasmic reticulum Ca^2+^-ATPase (SERCA) channels, resulting in a clearly distinguishable depletion of intracellular Ca^2+^ stores in the early phase and subsequent activation and opening of plasma membrane calcium channels in the late phase. Cell stimulation with thapsigargin caused significantly lower Ca^2+^-efflux from the ER store in the early and late phase of the measurement in THP-1 RTN1A^+^ cells (Fig 5J). The results show that RTN1A has an inhibitory effect on the Ca^2+^ efflux from ER in comparison to THP-1 wt cells and most likely results in a decreased Ca^2+^ influx through the plasma membrane. Together, our data suggest that RTN1A expression may induce a feedback regulatory mechanism to counteract elevated cell adhesion and further highlights a potential role of RTN1A in fine-tuning cell activation and adhesion. These results comply with a previous finding of the inhibitory effect of RTN1A on calcium release and therefor calcium flux in nerve cells^55^ and suggest its involvement in myeloid cell adhesion via regulation of integrin activation^56^.

### Stimulation of TLR1/2, TLR2 and TLR7 significantly diminishes RTN1A expression levels in eLCs and induce rLC clustering

We next explored whether a relation exists between RTN1A and the activation of LCs via TLR agonists, thus mimicking inflammatory conditions in *ex vivo* human skin. This was of great interest as it may reflect the everyday life of human skin which is constantly exposed to myriad environmental assailants. Incubation of epidermal explants with selected extra- and intracellular TLR agonists and subsequent analysis of eLCs by flow cytometry (Fig 6A and S. Fig 3A) revealed that both, the percentage of RTN1A^+^ eLCs and RTN1A expression intensity in eLCs were significantly diminished after activation with the agonists TLR1/2 (Pam3CSK4), TLR2 (listeria monocytogenes), and TLR7 [imiquimod, polyadenylic:polyuridylic acid (poly(A:U))], but not to agonists of TLR2/6 (mycoplasma salivarium) and TLR3 [low and high molecular weight polyinosinic:polycytidylic acid (LMW and HMW poly(I:C))] (Fig 6B-D). Of note, the downregulation of RTN1A in eLCs was transient, since we observed a tendency for recovery of RTN1A protein expression upon 48 h of cultivation (Fig 6E). Next, we examined whether eLCs diminish RTN1A expression during the activation process. In cultures with untreated epidermal sheets, we observed a small percentage of activated eLCs (S. Fig 3B,C). This is in line with previous observations that some rLCs can be activated after enzymatic separation and cultivation^57^. Epidermal sheets cultured with TLR1/2, TLR2/6 and TLR3/7 agonists and analysis of pre-gated RTN1A^+^ eLCs revealed an upregulation of CD83 and CD86 in comparison to the untreated control (S. Fig 3B,C). Of note, TLR1/2 stimulation significantly enhanced the percentage of activated CD83^+^CD86^+^ eLCs (S. Fig 3B,C). These data provide evidence that the decreased RTN1A expression correlates with the activation status of LCs and implies an active communication between TLRs and RTN1A. Next, the expression and distribution of RTN1A, CD86 and CD83 were examined in/on rLCs in epidermal sheets after cultivation with selected extra- and intracellular TLR agonists. Stimulation with TLR1/2, TLR2 and TLR7 agonists induced the formation of rLC clusters and were repeatedly detected in the anfractuous epidermal areas (Fig 7A, inserts). The big and small rLC clusters exhibited diminished dendrites and a roundish morphology (Fig 7B). Furthermore, we observed low co-localization between activation markers (cell membrane) and RTN1A (ER) in the single, activated and dendritic rLCs (Fig 7B). In contrast, upon stimulation of epidermal sheets with TLR2/6 and TLR3 [LMW and HMW p(I:C)] agonists, the morphology of rLCs was unaltered and comparable to the untreated control. The expression intensity of RTN1A in rLCs was slightly but not significantly reduced upon stimulation with all TLR agonists in comparison to the untreated control (Fig 7C). Significant upregulation of CD83/CD86 was observed after TLR7 stimulation (Fig 7D). Further analysis of clusters formed by rLCs upon TLR1/2 and TLR7 stimulation of epidermal sheets revealed that they were not proliferating (S.Fig 3D). They simultaneously acquired a migratory phenotype and upregulated either MMP-9 or CCR7 or co-expressed both markers (Fig 7E, yellow: MMP-9/CCR7, red: MMP-9, and turquoise: CCR7). Stimulation of TLRs activates signaling cascades inducing proinflammatory response such as secretion of cyto- and chemokines^38^. Accordingly, we have assessed epidermal sheet culture supernatants and detected low IL-6 levels with exception of a significant increase after stimulation with TLR3/7 poly(A:U). IL-8 and TNF-α concentrations were significantly elevated after stimulation with TLR3-HMW, and TLR3-LMW agonists, respectively. Of note, MCP-1 levels were significantly reduced after stimulation with TLR2 and TLR3/7 poly(A:U) agonists in comparison to untreated control (Fig 7F). In contrast, IL-10, IL-23 concentrations were slightly but not significantly elevated in comparison to IL-6 and IL-8 (S. Fig 3E).

**Fig 6.**
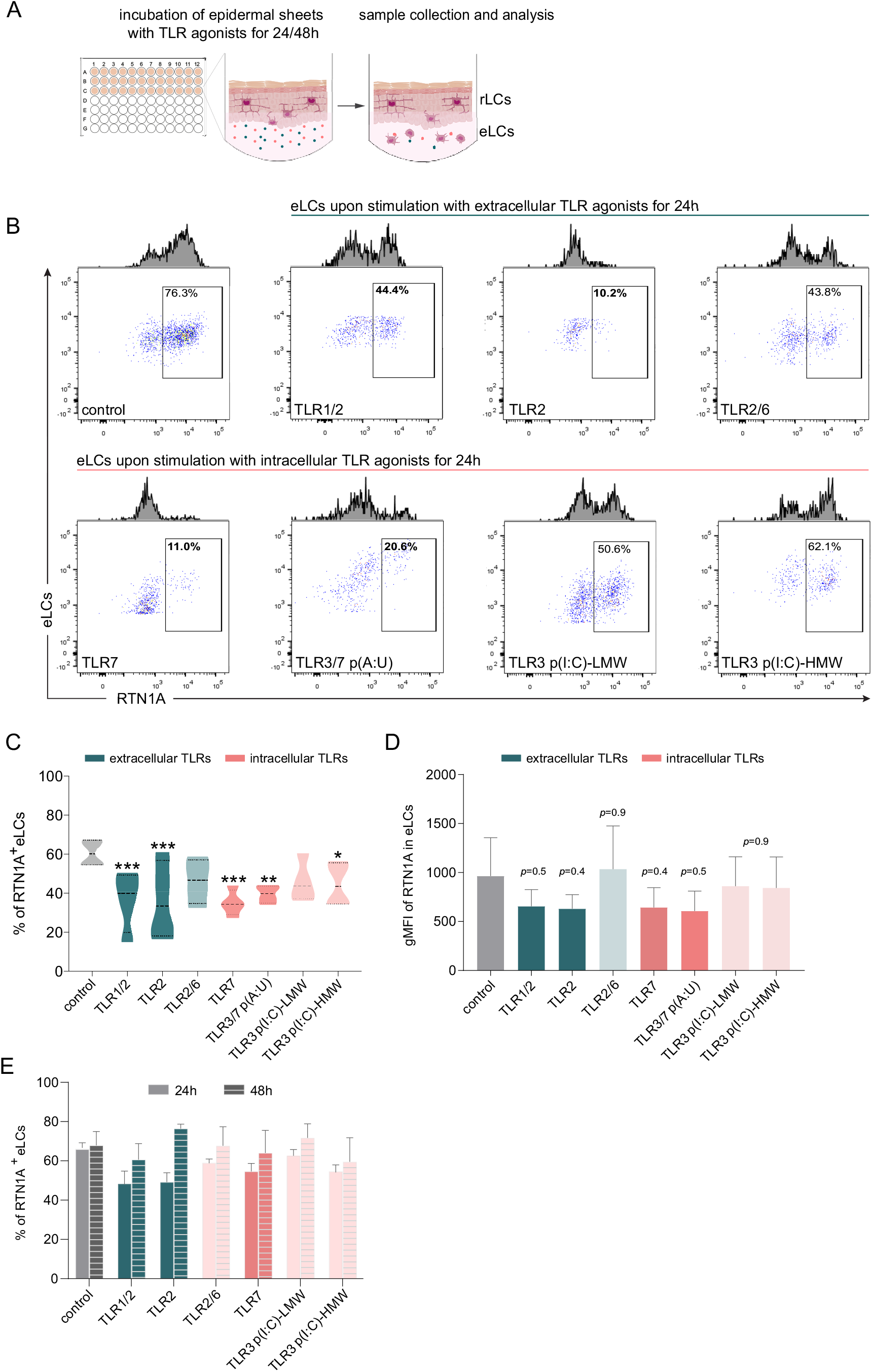
Emigrated LCs (eLCs) significantly decrease RTN1A expression upon stimulation with TLR1/2, TLR2 and TLR7 agonists. A. Graphical presentation for the stimulation of epidermal sheets with TLR agonists and analysis. B. Representative FACS blots of RTN1A expression in pre-gated eLCs upon stimulation of epidermal sheets with TLR1/2 (Pam3CSK4), TLR2 (listeria monocytogenes), TLR2/6 (mycoplasma salivarium), TLR7 (imiquimod), TLR3/7 [polyadenylic:polyuridylic acid (poly(A:U)], and TLR3 [low and high molecular weight polyinosinic:polycytidylic acid (LMW and HMW poly(I:C))] for 24 h. C, D. The percentage and geometric mean fluorescence intensity (gMFI) of RTN1A in eLCs are shown as standard error of the mean (SEM) of triplicates, and analyzed using two-way ANOVA with Tukey’s multiple-comparison test. \**p* ≤ 0.05, \*\**p* ≤ 0.01, \*\*\**p* ≤ 0.001. Some *p* values were not significant (ns), yet indicative of a trend for the reduction of RTN1A expression intensity. E. Recovery of the RTN1A protein expression (% of RTN1A^+^ eLCs) after 48 h of cultivation with indicated TLR agonists. Data shown represent mean ± SEM of triplicates from 3 donors.

**Fig 7.**
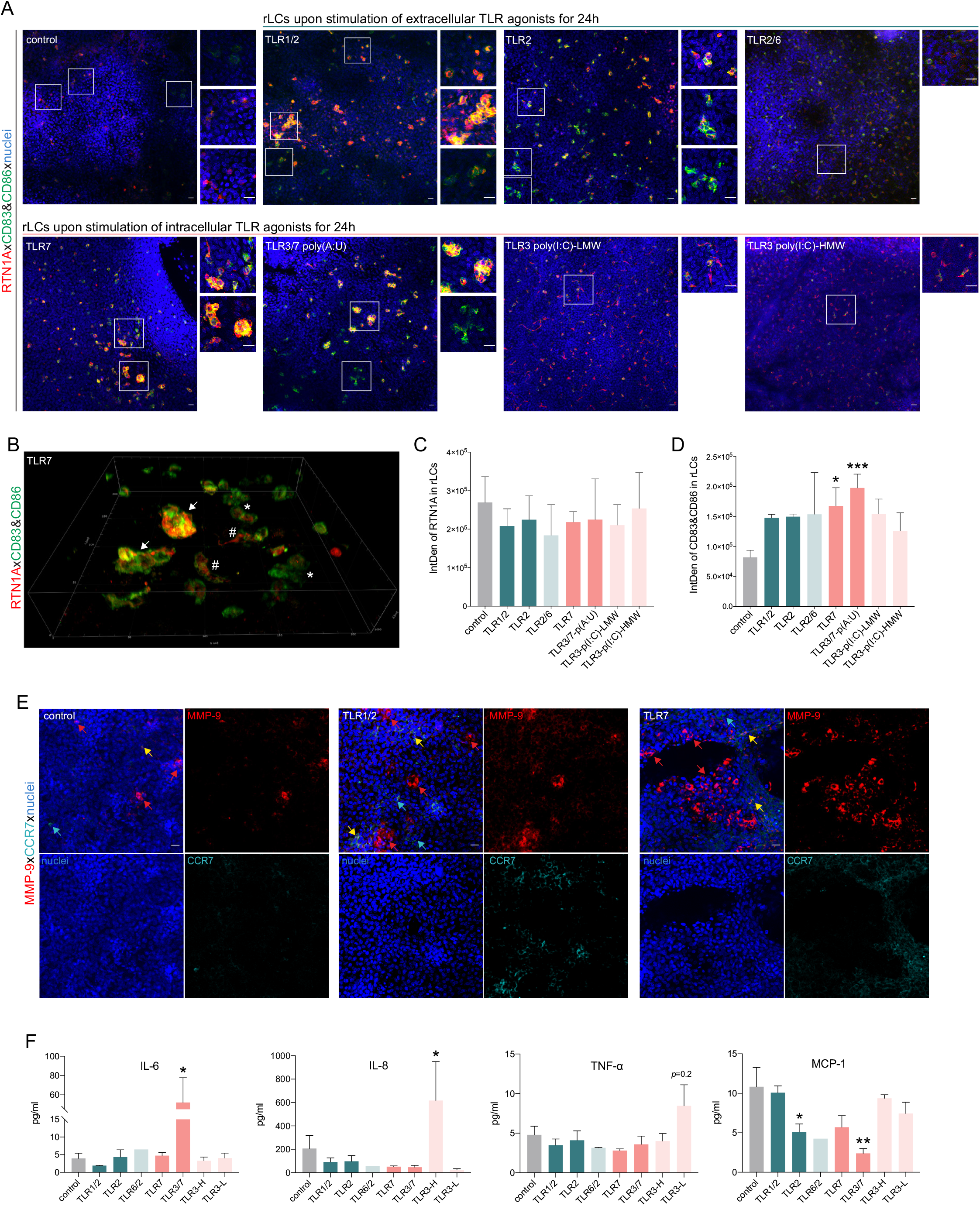
Stimulation of epidermal sheets with TLR1/2, TLR2 and TLR7 agonists initiates cluster formation of resident Langerhans cells (rLCs). A. Representative immunofluorescence (IF) images of untreated (control) and indicated TLR-treated epidermal sheets upon 24 h of culture stained with RTN1A (red), CD83, CD86 (green) and DAPI (blue). n=5, scale bar: 20 μm. B. 3D projection of rLCs in an epidermal sheet after incubation with a TLR7 agonist for 24 h (A). rLCs form big (arrow) and small (asterisk) clusters yet were also visible as single activated dendritic rLCs (hashtag). C, D. Expression intensity of indicated markers in rLCs upon culture and treatment. Data are shown as standard error of the mean (SEM) representing 4 fields of view of 5 donors, analysed with two-way ANOVA Tukey’s multiple comparison test. *\*p ≤* 0.05,*\*\*\*p ≤* 0.001E. E. Representative IF images of untreated (control) and TLR-stimulated epidermal sheets stained for MMP-9, CCR-7 and DAPI. n=2, scale bar: 20 μm F. Inflammatory cyto- and chemokine concentrations in supernatants of epidermal sheet cultures after 24 h, measured in duplicates with LEGENDplex bead array. Data are shown as SEM of 4 donors and analyzed using two-way ANOVA with Durrett’s multiple comparison test). \**p ≤* 0.05, **p *≤* 0.01.

## Discussion

In this study, we investigated the involvement of RTN1A in structural remodeling and effects of immune responses against pathogens by stimulation of TLRs in rLCs and eLCs in human epidermis *ex vivo*. We showed that local attenuation of RTN1A functionality with an N-terminus-targeting ab caused significant changes in the morphology of rLCs, leading to a roundish cell body with reduced dendrite length and dendrite distribution. Labelling of rLCs in human skin *ex vivo*^58^ and in mouse skin *in vivo*^59^ has been successfully employed before, yet in this study we targeted an intracellular protein. These observations led us to conclude that RTN1A is promoting elongation of dendrites in rLCs. Similar results with involvement of RTN1 family members in dendrite formation have been reported for Purkinje cells in mice^62^ and cutaneous nerves^25^ in developing mouse skin. Of note, other RTN family member such as RTN4 isoforms are vice versa involved in inhibition of neurite regeneration and elongation as well as impairment in sphingomyelin processing^30,60,31,61^. Our further observations on effects and consequences of RTN1A expression in THP-1 cells, such as significantly smaller cell size and a denser intermediate filament constellation, and in contrary in differentiated adherent THP-1 RTN1A^+^ Mφs the increased capacity to form long cell protrusions and cell body, support our hypothesis on the crucial role of RTN1A in cytoskeleton dynamics. Moreover, the excessive colocalization with vimentin (Fig 1D, Fig 4A, Fig 5E), but less with F-actin, could indicate an interaction between ER tubules and intermediate filaments, thereby facilitating changes in cell morphology, when RTN1A is inhibited or overexpressed. Cytoskeletal components such as type III intermediate filaments (e.g. vimentin)^63^ were shown to i) promote cell adhesion through governing integrin functions^64,65^, ii) stabilize microtubule dynamics^66^, and iii) endorse cell migration^67^ by contact-dependent cell stiffening^68^. Alterations in the adhesion molecule profile after inhibition of RTN1A in rLCs and THP-1 RTN1A^+^ cells further support our assumption that RTN1A is crucial for structural cell dynamics. Based on our findings we suggest that RTN1A could interact with intermediate filaments by acting as GTPase adaptor molecule, which will be investigated in the future.

Our discovery that RTN1A in THP-1 cells significantly reduces Ca^2+^ flux after stimulation with ionomycin is in line with results showing that RTN1A impaired the Ca^2+^ flux in Purkinje cells by inhibition of the RYR calcium channel^55^ and SERCA2b^69^. Apparently, RTN1A can inhibit calcium flux in cells with an extensive dendrite network such as Purkinje cells, THP-1 RTN1A^+^ cells (this study) and presumably LCs. Calcium signaling leads to cell activation^70^, and lower calcium flux is supposed to protect the cell from unnecessary activation and/or maintain their steady state within the tissue^71,72^. Indeed, we found, that application of the inhibitory RTN1A ab to epidermal sheet cultures prevented an upregulation of CD86 and consequently activation of LCs. Human LCs express a particular repertoire of extracellular TLRs such as TLR1/2, TLR2^73^ for detection of bacterial lipids, and intracellular TLR7^16^ but not TLR3^74^ for recognition of viral RNA. While assessing the effects of extra- and intracellular TLR-stimulation on the expression and distribution of RTN1A in rLCs in epidermal sheets, we found that stimulation with TLR1/2, TLR2 and TLR7 agonists but not with TLR2/6 and TLR3 agonists significantly diminished the percentage and expression intensity of RTN1A in eLCs in culture wells (Fig 4C,D). As activation of TLR1/2, TLR2 and TLR7 leads to the generation of endolysosoms^35,75^, we postulate that the temporary downregulation of RTN1A in activated eLCs was due to the recruitment of RTN1A into endosomes. Essentially, it was shown that RTN3L (RTN3A), another member of the RTN family, is involved in endosome maturation^76^ and in autophagy-induced fragmentation of ER tubules^77^. Furthermore, the morphological changes in activated eLCs could cause repression of the ER-tubular network during the collision of late endosomes or lysosomes carried along microtubules^78^, despite of the fact that in unstimulated eLCs RTN1A expression is unchanged (Fig 6A,B). Moreover, stimulation of TLR1/2, TLR2 and TLR7, which impacts RTN1A expression, induced also cluster formation by rLCs within epidermal sheets (Fig. 5C). The clustering rLCs acquired an activated and migratory phenotype^79,80^. To our knowledge, this behavior of rLCs in human epidermis has not been observed before. Previously, it was reported that dermal plasmacytoid DCs in mice can cluster in the dermis after topical stimulation with the TLR7 agonist imiquimod regulating antiviral response, but not epidermal LCs, which after several days of treatment appeared to be larger but not clustered^81^. A similar event was described for dermal CD11c^+^ DCs but not for LCs in a contact dermatitis mouse model using hapten sensitization^82^. Recently, it was demonstrated that in the dermis of atopic dermatitis patients perivascular leukocyte clusters can be infiltrated by other antigen presenting cells to regulate T cell activation^83^. As we have investigated the LC activation process in the early stages of inflammation-like conditions, minor levels of signature inflammatory cytokines produced usually by LCs and specific for the response to applied stimuli such as IL-1α, IL-1β, IL-6, IL-8, TNF-α^38^ or IFN-γ^84^ were measured.

In conclusion, we demonstrated the importance of RTN1A in LC intra-tissue dynamics, suggesting its strong involvement in maintaining homeostasis in rLCs. Moreover, we demonstrate a close functional relation between RTN1A and particular TLRs at the active stage of LCs, which could have a protective role in the maintenance of tissue homeostasis.

## Methods

### Ethics statement

Abdominal skin samples from anonymous healthy donors (female/male, age range: 27-44 years) were obtained during plastic surgery procedures. The study was approved by the local ethics committee of the Medical University of Vienna and conducted in accordance with the principles of the Declaration of Helsinki. Written informed consent was obtained from all participants.

### Processing of human skin

Experiments were performed within 1-2 hours (h) after surgery. Skin was dermatomed (Aesculap) to 600 µm thickness, then incubated with 1.2 U/ml dispase II (Roche Diagnostics) in Roswell Park Memorial Institute 1640 medium (RPMI; Gibco, Thermo Fisher Scientific) with 1% penicillin/streptomycin (P/S; Gibco, Thermo Fisher Scientific) overnight (ON) at 4°C. After washing with phosphate buffered saline (PBS; Gibco, Thermo Fisher Scientific), epidermis was separated from the dermis. Epidermal punches with a diameter of 6 mm were obtained and used for the following experiments. The rLCs were isolated and purified as described previously^74^.

### Cultivation of epidermal explants with TLR agonists and staining

Epidermal punch biopsies (6 mm diameter) in triplicates for each condition were floated on RPMI medium supplemented with 10% fetal bovine serum (FBS, Gibco), 1% P/S and human TLR agonists (InvitroGen) in 96-well round bottom plates.. The following TLR agonists were used: TLR1/2 (Pam3CSK4×3HCl; 1 µg/ml), TLR2 [heat-killed *listeria monocytogenes* (HKLM); 10^8^ cells/ml], TLR3/7 [(poly(A:U); 25 µg/ml], TLR3 [LMW and HMW poly(I:C); both at 0.8 µg/ml], TLR6/2 (mycoplasma salivarium, FSL-1, Pam2CGDPKHPKSF; 2.5 µg/ml) and TLR7 (imiquimod; 2.5 µg/ml). After 24 and 48 h, epidermal sheets were collected, fixed with acetone (Merck) for 10 minutes (min) at room temperature (RT), incubated with FITC-conjugated α-CD83 and α-CD86 abs as well as with a primary mouse α-RTN1A ab (mon162, abcam) ON at 4°C. All abs were diluted in 2% bovine serum albumin (BSA; Gibco, Thermo Fisher Scientific) in PBS. Subsequently, sheets were incubated with a AF647-labeled α-mouse secondary ab (Thermo Fisher Scientific) for 1 h at RT and counterstained with 4′,6-diamidino-2-phenylindole (DAPI; Sigma-Aldrich) for 1 min. After PBS washing, sheets were mounted (ProLong™ Gold antifade reagent, Invitrogen) with the *stratum corneum* facing the slide and imaged. At the same time points cultivation medium was harvested and stored at -80°C for further analyses and eLCs were processed and analyzed as described below.

### Cultivation of epidermal explants with an α-RTN1A ab

Epidermal punch biopsies (6 mm diameter) in triplicates were floated on medium containing either a mouse α-RTN1A ab (mon162, abcam) or the respective isotype control ab (IgG1, abcam) (5 µg/ml/each) in 96 well round bottom plates at 37°C, 5% CO_2_. Sheets were collected at 3, 6, and 24 h, fixed with acetone and stained with a secondary α-mouse cross-absorbed F(ab’) ab fragment conjugated with AF546 (Thermo Fisher Scientific) and a FITC-labelled α-CD207 (Dendritics) to identify LCs. In some experiments, 24 h cultured epidermal sheets were stained with a FITC-labelled α-vimentin, primary rat α-CD207 (Sigma Aldrich) and mouse α-RTN1A abs (mon162, abcam) ON at 4°C, followed by α-mouse and α-rat secondary abs and counterstaining with DAPI. 3D-projections have been created with ImarisViewer (v.9.8). The contrast and brightness of representative images (Fig 1 B-C) remained unaltered, to highlight the detection level of the abs taken up by rLCs. All abs and reagents used in this study are listed in **Error! Reference source not found**.

### Generation of an RTN1A expressing THP-1 cell line

Human RTN1A (NM_0211369) was gene synthesized (Eurofins), cloned into the lentiviral expression vector pHR-SIN-BX-IRES-Emerald^85^ and expressed in the THP-1 cell line. Following puromycin selection, stable RTN1A protein expression was tested by flow cytometry (α-RTN1A-APC labelled ab, mon161, Novus Biologicals, Biotechne).

### Differentiation of THP-1 wt and THP-1 RTN1A ^+^cells towards Mφs

THP-1 wt and THP-1 RTN1A^+^ cells were seeded on cover slips in a 24 well plate (2.5×10^4^ cells/well) and polarized towards Mφs for 72 h with 50 ng/ml phorbol 12-myristate 13-acetate (Abcam) in RPMI medium and supplements as described previously^86^.

### Immunofluorescence staining of THP-1 cells and THP-1 Mφs

THP-1 wt and THP-1 RTN1A^+^ cells (both seeded at 2×10^4^) on adhesion slides (Marienfeld) were fixed with acetone for 10 min at RT, stained with α-vimentin and α-RTN1A abs and mounted. For colocalization assays, THP-1 RTN1A^+^ cells were seeded in 8-well chamber slides (2×10^4^/well, ibidi), coated with 0.1 ng/ml of fibronectin (PromoCell). After 24 h the cells were washed, fixed with 4% formaldehyde (SAV Liquid Production) for 10 min at RT and permeabilized with 0.1% Triton™ X-100 (Sigma Aldrich) in PBS for 10 min at 4°C. These samples were stained additionally with phalloidin-AF647 probe (F-actin, Invitrogen) for 1 h at RT. Mφs differentiated on cover slips, were fixed and processed as described above for THP-1 RTN1A^+^ cells on coated slides.

### Microscopy and image analysis

A confocal laser scanning microscope (Olympus, FLUOVIEW-FV 3000, equipped with OBIS lasers: 405, 488, 561, 640 nm and 20x, 40x or 60x UPlanXApo objectives) and Olympus FV31S-SW software were used in this study. Images were acquired with 20x objective as Z-stack from 4 fields of view (FOV) per epidermal sheet from 4 different donors and analyzed using ImageJ Fiji software^87^. The measurement of the integrated density from the region of interest (ROI) was based on Z-projections with max intensity of manually thresholded images (analogues parameters were used for all analyzed images). Between 100-200 ROIs (rLCs) were analysed per 4 FOVs.

### Evaluation of the morphology and dendricity of rLCs and THP-1 Mφs

The enumeration of roundish (none or 1 dendrite) and dendritic (2 or more dendrites) rLCs per 0.04 mm^3^ in epidermal sheets was based on vimentin staining and performed using ImageJ Fiji. 60-300 cells were analysed per FOV. The average length of rLC dendrites and the distance of dendrites from the middle of the cell body was analysed and quantified using simple neurite tracer (SNT) plugin in ImageJ Fiji^88^. 60x objective was used for representative images in Figure 2A. The length of cell protrusions in Mp0 was also quantified using SNT (10-40 cells/FOV from 4 FOVs).

### Analysis of THP-1 and THP-1 Mφ cell areas and colocalization of RTN1A with cytoskeleton structures

To estimate the full size of the cell body of vimentin- and F-actin-stained Mφs on adhesion slides, vimentin and F-actin channels were merged, thresholded and the ROI area analyzed and quantified using ImageJ Fiji. To assess colocalization of RTN1A with vimentin and F-actin in THP-1 RTN1A^+^ cells and THP-1 Mφs, we used Manders’ coefficient analysis with RTN1A as M1 and vimentin/F-actin as M2. Single Z-stack slices from the bottom, middle and the top of 10 cells/FOV from 4 FOV were analyzed.

### Flow cytometry analysis of eLCs

eLCs were collected, stained and analysed with flow cytometry as described previously^25^. Briefly, eLCs were washed with PBS (Gibco, Thermo Fisher Scientific), stained with fixable viability dye and an ab cocktail for LC surface markers [PE-conjugated CD207 (Beckman Coulter), CD1a, HLA-DR (BD Pharmingen)], activation [FITC-labelled CD83 (BD Pharmingen)] and costimulatory markers [CD86 abs (BD Pharmingen)]. Next, cells were fixed, permeabilized and stained intracellularly with an APC-conjugated α-RTN1A ab (mon161, Novus Biologicals, Biotechne). The samples were acquired using FACS Verse (BD Biosciences) and BD Suite software (v1.0.5.3841, BD Biosciences). The gating strategy included discrimination of doublets and dead cells. Data was analyzed using the FlowJo software (v10.0.7r2, BD Biosciences). Mean percentages of positive cells and gMFI values from triplicates including 5 donors were analyzed using GraphPad Prism (v8.0.1). eLCs after ab uptake were collected and analyzed for the presence of primary abs by staining of viable eLCs with secondary α-mouse cross-absorbed F(ab’) antibody fragment conjugated with AF546 (Thermofisher Scientific), followed by CD1a (BD Pharmingen), CD207 (Beckman Coulter), CCR7 (Miltenyi Biotec) and CD86 (BD Pharmingen) staining and flow cytometry analyses.

### Measurement of adhesion molecules, cytokines and chemokines in epidermal explant and cell culture supernatants

LEGENDplexTM Human Adhesion Molecule Panel (13-plex) w/VbP LEGENDplex™ (Biolegend) and human inflammatory panel 1 (13-plex) w/VbP (Biolegend) was used to analyze supernatants from epidermal sheet and cell culture. The assay was carried out according to manufacturer’s instruction. Data analysis was performed using LegendPlex v8.0 software (BioLegend).

### Comparative evaluation of cell aggregate formation by THP-1 wt and THP-1 RTN1A^+^cells

Images of THP-1 wt and THP-1 RTN1A^+^ cells in cultures were taken with the ZOE fluorescence cell imager (Bio-Rad). and the numbers of cellular aggregates per 0.7 mm^2^ from 4 FOV have been quantified using ImageJ Fiji (Fig 5G). For enumeration of small and big clusters an average area of 312.5 µm^2^ and 637.6 µm^2^ were chosen, respectively (S. Fig 1 C).

### Measurement of cell proliferation and cell number

CellTrace™ CFSE cell proliferation dye (Invitrogen) was used according to manufacturer’s instructions. Briefly, THP-1 wt and RTN1A^+^ cells were incubated with CFSE (5 mM) for 20 min at 37°C, washed and seeded in a 24 well plate (2×10^4^ cells for each condition). Cells were collected at 0, 24, 48 and 72 h and acquired with FACS Verse. In parallel, the cell number was assessed at the same time points with trypan blue solution (Sigma) and Neubauer chamber (BRAND).

### Measurement of calcium flux in THP-1 wt and THP-1 RTN1A^+^cells

Ratiometric calcium flux experiments with Fura Red (Invitrogen) were performed similar to a previously described method ^52^. Briefly, 1×10^6^ THP-1 wt and RTN1A^+^ cells were washed, resuspended in 100 μl medium containing 1 μM Fura Red and incubated 30 min at 37°C. Cells were washed once with medium, resuspended in 1 ml medium and incubated for another 30 min at 37°C. Thereafter, cells were rested on ice until data acquisition at a FACSAria III flow cytometer (BD Bioscience). For the measurement of intracellular calcium flux, 300 μl Fura Red-loaded cells were transferred to FACS tube, pre-warmed for 5 min at 37°C and the baseline response was recorded for 30 seconds. After adding with 1 μg/ml ionomycin (InvivoGen) or thapsigargin (Invitrogen), cell responses were recorded for 5 min to analyze changes in calcium mobilization. Fura Red was excited using a violet laser (405 nm) and a green laser (561 nm) and changes in emission were detected with a 635LP, 660/20 BP and a 655LP, 795/40 BP filter set, respectively. The ratiometric ‘Fura Red Ratio’ over time was calculated using the Kinetics tool in FlowJo software version 9.3.3 (Tree Star Inc.) as follows:

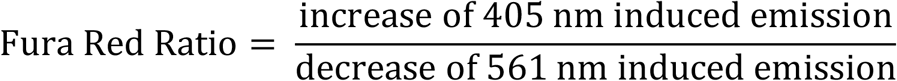

### Statistical analysis

Statistical analysis of the data has been performed using GraphPad Prism (v8.0.1) software. The number of technical and biological replicates have been implicated in respective method sections and figure legends. The statistical tests were adapted to the experimental design: for comparison of two samples (student t test), for higher number of samples with replicates (two-way ANOVA with Tukey’s or Durrett’s multiple comparison test). In some figures *p* values were displayed to indicate a tendency, despite lacking significance. Asterisks indicate significant p values; ns-not significant, p ≥ 0.05, *p ≤ 0.05, **p ≤ 0.01, ***p ≤ 0.001 and ****p ≤ 0.0001.

---

**Supplementary Fig 1.**
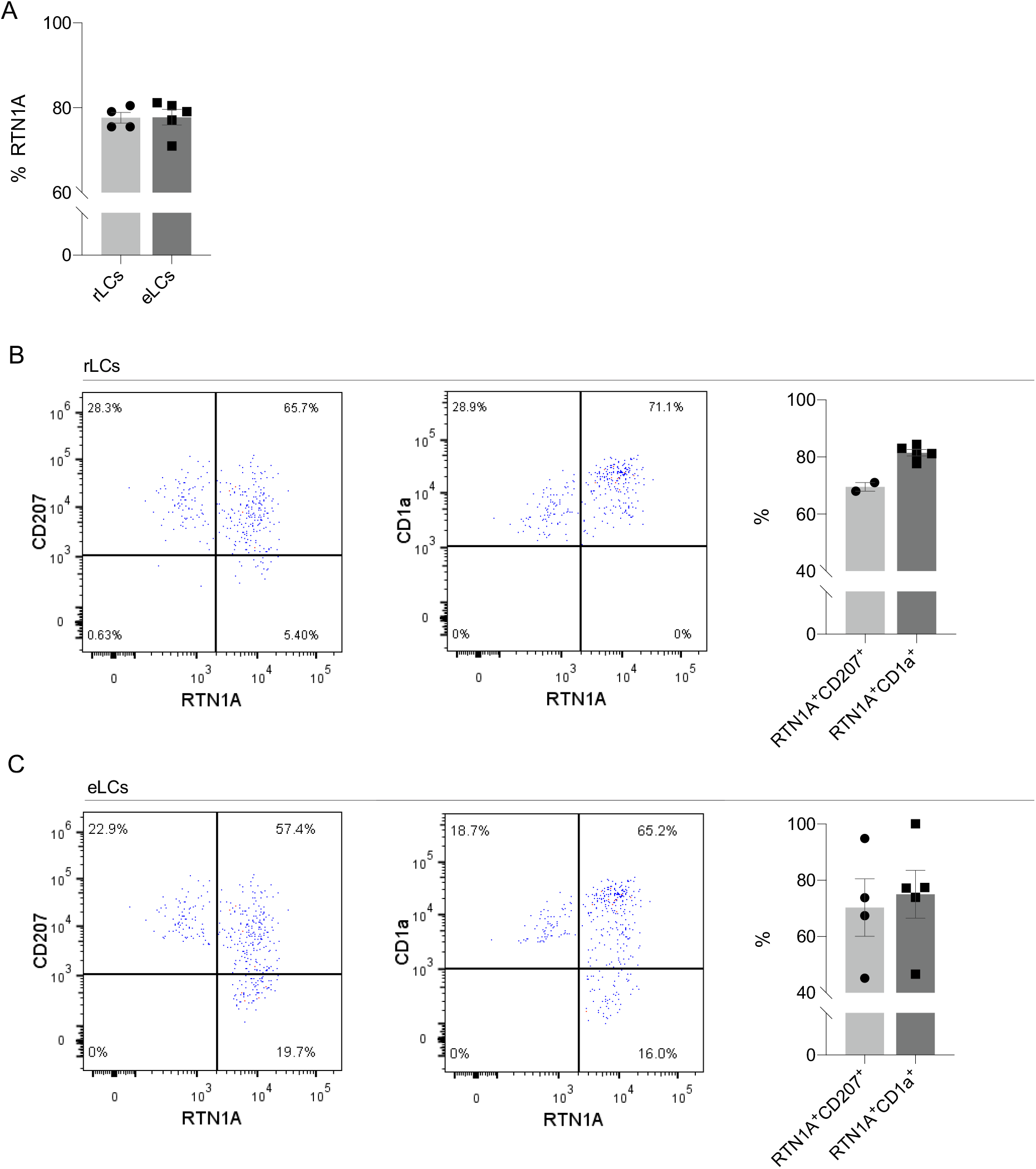
A.Percentage of total RTN1A^+^ resident Langerhans cells (rLCs) and emigrated LCs (eLCs), analysed with flow cytometry. Data shown as standard error of the mean (SEM), each dot represents one donor. rLCs: n=4, eLCs= n=5. B, C. Frequency of RTN1A expression in LCs. Representative dot blots showing flow cytometry analysis of (B) freshly isolated rLCs and (C) eLCs, stained for CD207, CD1a, and RTN1A. Quantification of the percentage of double-positive LCs s shown. Data shown as standard error of the mean (SEM), each dot represents one donor. CD207: n=2-4, CD1a: n=5

**Supplementary Fig 2.**
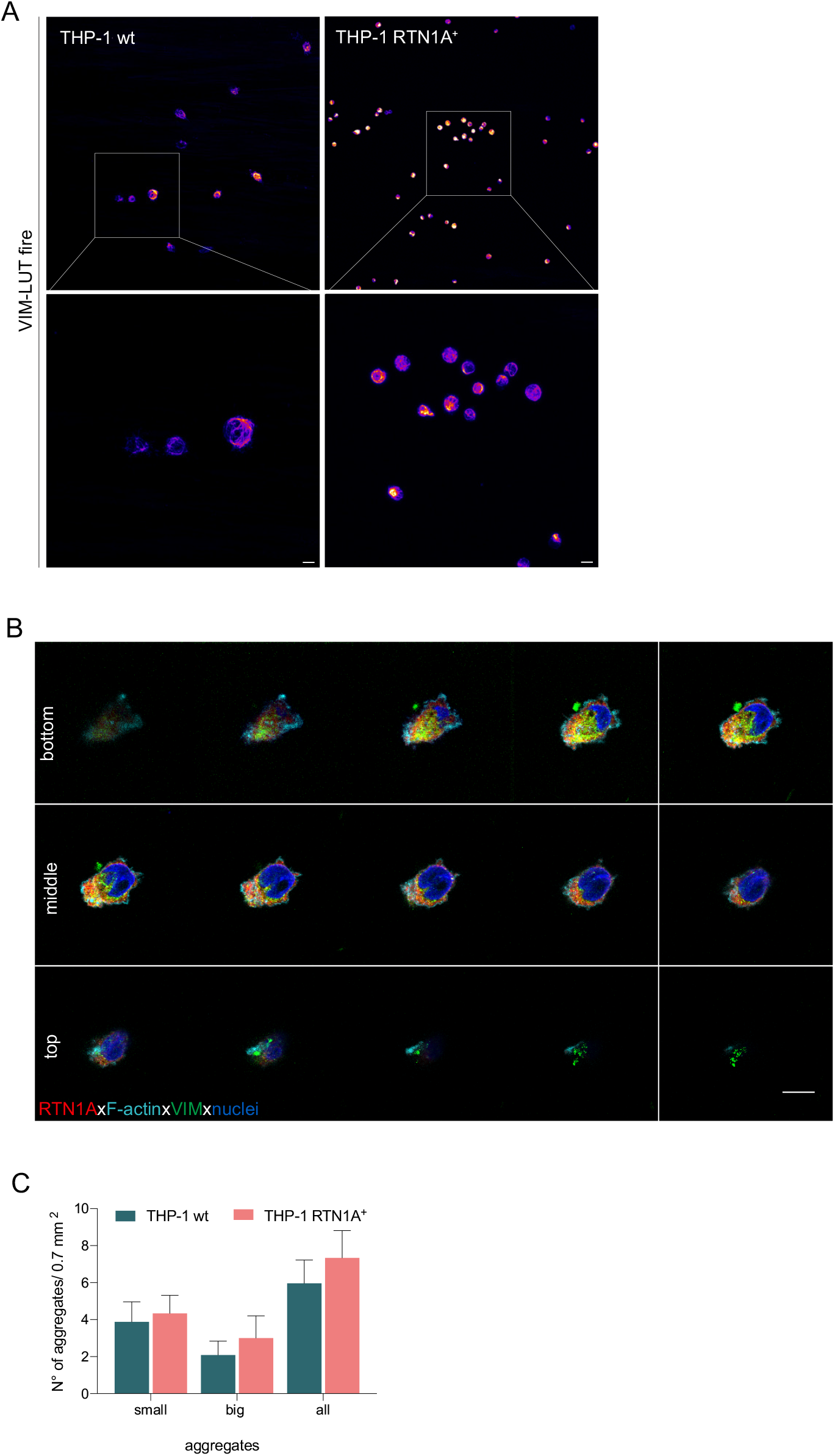
A.Representative immunofluorescence (IF) images of THP-1 wt and THP-1 RTN1A^+^ cells on adhesion slides stained with vimentin (VIM) and demonstrated as LUT-fire. THP-1 RTN1A^+^ cells display markedly more condensed intermediate filament structures in comparison to THP-1 wt cells. Scale bar: 10 μm. B. Representative IF image of THP-1 RTN1A^+^ cells stained for RTN1A, F-actin, VIM and nuclei. Image is shown as montage of Z-stack slices. C. Enumeration of small and big THP-1 wt and THP-1 RTN1A^+^ cell aggregates. n=6

**Supplementary Fig 3.**
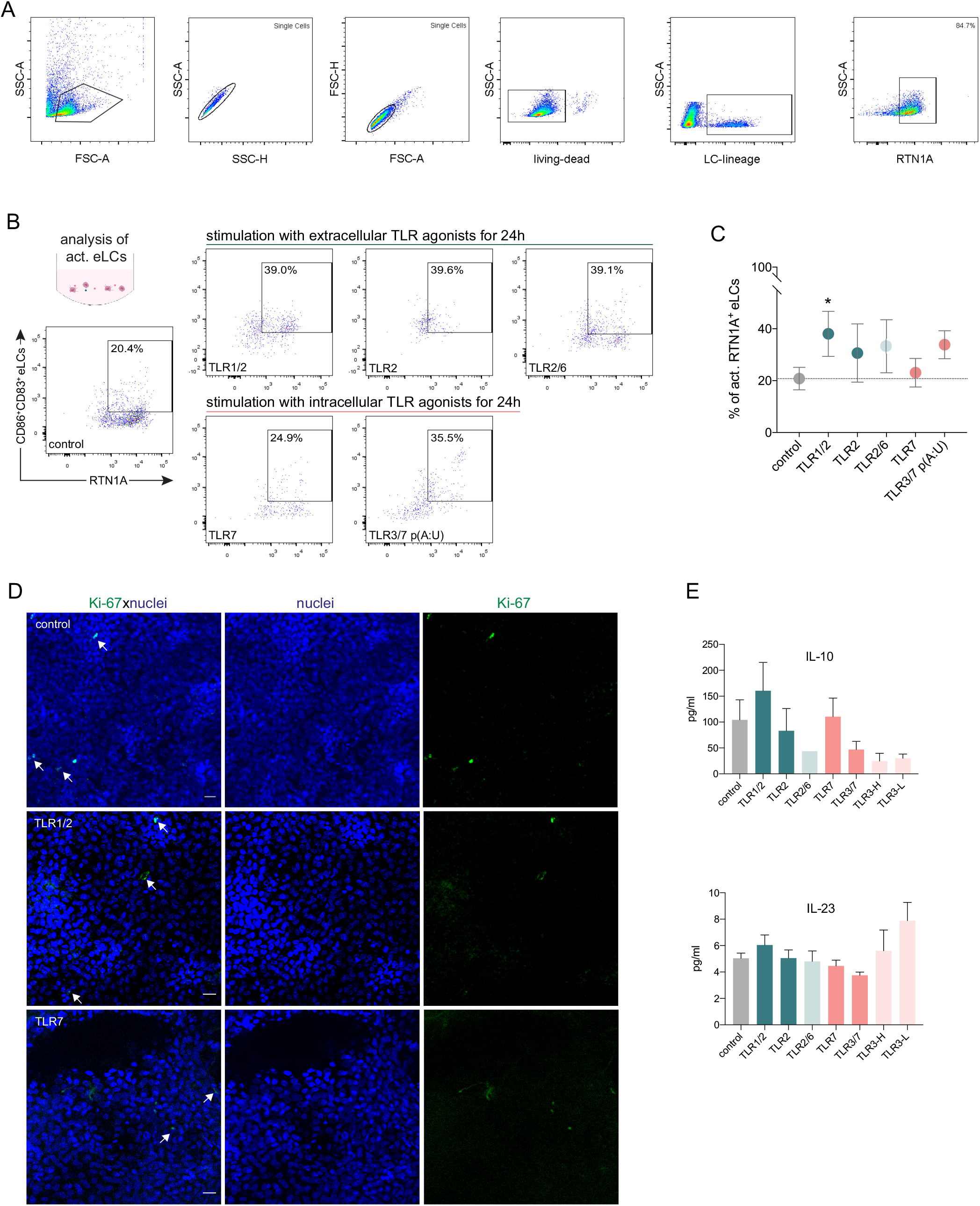
A.Gating strategy for emigrated Langerhans cells (eLCs) as analysed with flow cytometry. B, C RTN1A distribution within TLR-activated eLCs and their quantification. Data shown represent mean ± standard error of the mean (SEM) of triplicates from 4 donors. Data set was analysed using two-way ANOVA, Dunnett’s multiple comparison test. \**p ≤* 0.05D. D. Resident Langerhans cells (rLCs) do not proliferate in clusters upon TLR stimulation. Representative immunofluorescence images of epidermal sheets after 24 h of cultivation only or treatment with TLR 1/2, and TLR7 agonists. Epidermal sheets were stained for Ki67 (green) and nuclei (blue). n=2, scale bar: 20 μm. E. Inflammatory cyto- and chemokines measured in supernatants of THP-1 wt and THP-1 RTN1A^+^ cell cultures with LEGENDplex bead array after 24 h of cultivation. Data are shown as SEM from 4 donors and analyzed using two-way ANOVA with Durrett’s multiple comparison test and were not significant.

## Author Contribution

Conceptualization: M.A.C, A.E.-B.; Visualization: M.A.C; Data analysis: M.A.C, K.P.; Investigation and Methodology: M.A.C., J.L. K.P, L.W.; Resources: C.S., J.L., P.S., K.P.; Graphics: M.A.C; Project administration and supervision: A.E.-B.; M.A.C. and A.E.-B. wrote the manuscript; K.P., J.L., P.S., L.W. and C.S. proofread and edited the manuscript. All authors read and approved the manuscript. The authors declare no competing interests.

## Funding

This work was supported by the Austrian Science Fund (P31485-B30) and DK W1248-B30.

## Acknowledgements

We are grateful to all patient-volunteers, who donated material for this study. We thank Christian Schneeberger (Department of Gynecology, Medical University of Vienna) for providing access to the BD FACSVerse instrument. We thank our collogues from Department of Dermatology, Medical University of Vienna for support and scientific feedback.

---

**Table 1.** Antibodies and reagents used in this study

| Antibodies | Source | Identifier | Application |
| --- | --- | --- | --- |
| Mouse monoclonal α-RTN1A (clone: mon162), unconjugated | abcam | ab9274 | IF staining, cultivation of epidermal sheets |
| Mouse monoclonal α-RTN1A (clone: mon162), unconjugated | Novus Biologicals, Biotechnne | NBP1-97677 | IF staining, cultivation of epidermal sheets |
| Mouse monoclonal α-RTN1A-APC (clone: mon161) | Novus Biologicals, Biotechnne | NBP1-97678AF647 | FACS analysis |
| Mouse monoclonal IgG1 [15-6E10A7] - Isotype control, unconjugated | abcam | ab170190 | IF staining, cultivation of epidermal sheets |
| Rat monoclonal α-CD207-FITC (clone 929F3.01) | Dendritics | Cat:DDX0362 | IF staining |
| Rabbit monoclonal α-CD207, unconjugated | Sigma Aldrich | HPA011216 | IF staining |
| Mouse α-human CD207-PE (clone DCGM4) | Beckman Coulter | PN IM3577 | FACS analysis |
| Mouse α-human HLA-DR-PE | BD Pharmingen | Cat. 347401 | FACS analysis |
| Mouse α-human CD1a-PE | BD Pharmingen | Cat. 555807 | FACS analysis |
| Mouse α-human CD83-FITC | BD Pharmingen | Cat. 556910 | FACS analysis, IF staining |
| Mouse α-human CD86-FITC | BD Pharmingen | Cat: 555657 | FACS analysis, IF staining |
| CCR7-APC | Miltenyi Biotec | 5171113456 | FACS analysis, IF staining |
| Recombinant α-Vimentin-FITC (clone REA409) | Miltenyi Biotec | 130-116-508 | IF staining |
| Recombinant FITC isotype control | Miltenyi Biotec | 130-113-449 | IF staining |
| Alexa Fluor™ 647 Phalloidin | Invitrogen™ | REF A-22287 | IF staining |
| F(ab')<sub>2</sub>-Goat anti-Rabbit IgG (H+L) Cross-Adsorbed Secondary Antibody, Alexa Fluor 488 | Life Technologies | REF A-11070 | IF staining |
| F(ab')<sub>2</sub>-Goat anti-Mouse IgG (H+L) Cross-Adsorbed Secondary Antibody, Alexa Fluor 546 | Invitrogen™, Thermofisher scientific | REF A-11018 | IF staining |
| Alexa Flour 647 F(ab')<sub>2</sub> fragment of goat α-rabbit IgG (H+L) | Invitrogen™, Thermofisher scientific | REF A-21246 | IF staining |

| Alexa Fluor 555 goat α-mouse IgG (H+L) | Invitrogen™, Thermofisher scientific | REF A-21422 | IF staining |
| --- | --- | --- | --- |
| Fixable Viability Dye eFluor™ 450 | eBioscience | cat#65-0863-18 | FACS analysis |
| DAPI (4',6-Diamidino-2-phenylindole dihydrochloride) | Sigma-Aldrich | cat#D9542 | IF staining |
| Reagents | Source | Identifier |  |
| Dispase II (neutral protease, grade II) | Roche Diagnostics | 04942078001 |  |
| human TLR kit | InvitroGen™ | Cat. tlr-kit1hw |  |
| ProLong™ Gold antifade reagent | InvitroGen™ | Cat. P36934 |  |
| IC Fixation Buffer | eBioscience | cat#00-8222-49 |  |
| Permeabilization Buffer (10X) | eBioscience | cat#00-8333-56 |  |
| Phorbol 12-myristate 13-acetate (PMA), PKC activator | Abcam | Ab120297 |  |
| Triton™ X-100 | Sigma-Aldrich | T9284 |  |
| Fura Red™, AM, cell permeant | Invitrogen™, Thermofisher scientific | F3020 |  |
| CellTrace™ CFSE Cell Proliferation Kit, for flow cytometry | Invitrogen™, Thermofisher scientific | C34554 |  |
| Thapsigargin | Invitrogen™, Thermofisher scientific | T7458 |  |
| Ionomycin- Calcium ionophore - NFAT Activator | InvivoGen | inh-ion |  |
| Fibronectin Solution Human | PromoCell | Cat.No. C-43060 |  |
| LEGENDplex™ Human inflammatory Panel 1 (13-plex) w/VbP | Biolegend | Cat. 740809 |  |
| LEGENDplex™ Human Adhesion Molecule Panel (13-plex) w/VbP | Biolegend | Cat. 740946 |  |
| Softwares |  |  |  |

GraphPad Prism v8.0.1
FlowJo v10.6.1
Fiji: ImageJ
ImarisViewer 9.8
Adobe illustrator CS7

## Notes

### Competing Interest Statement

The authors have declared no competing interest.

